# Trait Stacking Simultaneously Enhances Provitamin A Carotenoid and Mineral Bioaccessibility in Biofortified *Sorghum bicolor*

**DOI:** 10.1101/2022.08.03.501587

**Authors:** Michael P. Dzakovich, Hawi Debelo, Marc C. Albertsen, Ping Che, Todd J. Jones, Marissa K. Simon, Zuo-Yu Zhao, Kimberly Glassman, Mario G. Ferruzzi

## Abstract

Vitamin A, iron, and zinc deficiencies are major nutritional inadequacies in sub-Saharan Africa and disproportionately affect women and children. Biotechnology strategies have been tested to individually improve provitamin A carotenoid or mineral content and/or bioaccessibility in staple crops including sorghum (*Sorghum bicolor*). However, concurrent carotenoid and mineral enhancement has not been thoroughly assessed and antagonism between these chemical classes has been reported. This work evaluated two genetically engineered constructs containing a suite of heterologous genes to increase carotenoid stability and pathway flux, as well as phytase to catabolize phytate and increase mineral bioaccessibility. Kernels from transformed sorghum events were processed into model porridges and evaluated for carotenoid and mineral content as well as bioaccessibility. Transgenic events produced markedly higher amounts of carotenoids (26.4 µg/g) compared to null segregants (4.2 µg/g) and wild-type control (Tx430; 3.7 µg/g). A 200 g serving of porridge made with these transgenic events represents a projected 53.7% of a 4– 8-year-old child’s vitamin A estimated average requirement. Phytase activation by pre-steeping flour resulted in significant phytate reduction (9.4 to 4.2 mg/g), altered the profile of inositol phosphate metabolites, and reduced molar ratios of phytate to iron (16.0 to 4.1); and zinc (19.0 to 4.9) in engineered material; suggesting improved mineral bioaccessibility. Improved phytate:mineral ratios did not significantly affect micellarization and bioaccessible provitamin A carotenoids were over 2300% greater in transgenic events compared to corresponding null segregants and wild-type controls. These data suggest that combinatorial approaches to enhance micronutrient content and bioaccessibility are feasible and warrant further assessment in human studies.

## 1.1 Introduction

Incidence of food insecurity is unusually high in sub-Saharan Africa where diets are dominated by carbohydrate-rich cereal grains (Fraval et al., 2019; Xu et al., 2021). As a result of food scarcity and dietary patterns in these regions, vitamin A, iron, and zinc deficiencies remain prominent (Black et al., 2008; Harika et al., 2017). Vitamin A must be obtained through the diet either as preformed retinol (through animal sources) or as provitamin A carotenoid precursors such as β-carotene (predominantly from plant-based sources). Chronic vitamin A deficiency can result in xerophthalmia; the leading cause of preventable blindness in children globally (Humphrey et al., 1992; Whitcher et al., 2001). Likewise, iron and zinc are required for normal growth and development and their deficiencies are associated with an increased risk of death from other diseases such as diarrhea (Trumbo et al., 2001; Fischer Walker et al., 2009). Considering social, economic, and environmental factors, food choice and availability are unlikely to markedly change in sub-Saharan Africa. Improvement of staple foods already predominant in the broader diet of at-risk populations is a viable strategy to ameliorate malnutrition.

Sorghum (*Sorghum bicolor* (L.) Moench) is a close relative of maize (*Zea mays*) and one of the most popular cereal grains consumed in sub-Saharan Africa (Peng et al., 1999). Sorghum is often processed into porridges (e.g. tô) by mixing flour with boiling water (Da et al., 1981).

However, sorghum and subsequent food products are generally low in micronutrients such as provitamin A carotenoids and marginally bioavailable iron and zinc (Kayodé et al., 2006; Kean et al., 2011). Low mineral contents are exacerbated by the presence of high levels of the antinutrient phytate; a major storage molecule for phosphorous located in the aleurone of cereal kernels that strongly chelates divalent metals like iron and zinc and reduces their bioavailability (Afify et al., 2011). Previous research using labeled mineral elements estimate that molar ratios of phytate to iron or zinc above 10-15 indicate poor bioavailability (Oberleas & Harland, 1981; Turnlund et al., 1984; Morris & Ellis, 1989; Saha et al., 1994). Depending on sorghum genetic background and environmental conditions, these ratios are overwhelmingly above these thresholds and vary between 6-55 for phytate to iron and 30-40 for phytate to zinc (Kayodé et al., 2006, 2007; Kruger et al., 2014). Given the low levels of carotenoids, high concentration of phytate, and dietary significance of the crop, sorghum is an ideal candidate for biofortification efforts to increase fat-soluble carotenoids or reduce the mineral-limiting antinutrient phytate.

Biofortification strategies seeking to simultaneously improve delivery of provitamin A and key shortfall minerals such as iron and zinc are rare. Improvements in carotenoid content, stability, and reducing antinutrients have been tested individually using enzymes such as homogentisate geranylgeranyl transferase (HGGT), phytoene synthase (PSY1), phytoene desaturase (CRTI), phytoene synthase (CRTB), and phytase (PhyA) among others (Khush et al., 2012; Lipkie et al., 2013; Che et al., 2016, 2019; Díaz-Gómez et al., 2017). Through biotechnological approaches, it is theoretically possible to address both provitamin A carotenoid and mineral content as well as minimizing factors that negatively impact bioavailability.

However, potential for antagonism between carotenoid bioavailability and divalent minerals has been reported in *in vitro* and *in vivo* studies (Biehler et al., 2011b; a; Corte-Real et al., 2016, 2017; Kopec et al., 2019). These effects are concentration dependent and their relationship in the context of sorghum biofortification remains unexplored, but critical to define.

To address this gap of knowledge, the present study utilized a coordinated approach by which tocotrienol and carotenoid biosynthetic genes (HGGT, CRTI, PSY1, and CRTB) as well as phytase (PhyA) were heterologously expressed to improve the bioaccessibility of both provitamin A carotenoids and mineral elements. Sorghum events were processed into model porridges and a three-stage *in vitro* digestion model was utilized to evaluate carotenoid and mineral bioaccessibility. Due to the intrinsically low mineral content of sorghum, we hypothesized that increased mineral release from phytate degradation would not significantly counteract the delivery of provitamin A carotenoids. Our results suggest that these transgenic sorghum events provide dramatically higher amounts of bioaccessible provitamin A carotenoids and exhibit altered phytate:iron/zinc molar ratios suggestive of enhanced mineral bioaccessibility.

## 1.2 Materials and Methods

### 1.2.1 Chemicals and reagents

Reagents sourced from Sigma Aldrich (Sigma Chemical Co., St. Louis, MO, USA) included mucin (M2378), α-amylase (A3176; 15.8 units/mg solid), pepsin (P7125), lipase (L3126), pancreatin (P7545), and bile (B8631). Urea (U15-500) and uric acid (A13346-14) were purchased from Fisher Scientific (Fisher Scientific, Waltham, MA, USA). The oral phase base solution contained potassium chloride, sodium phosphate, sodium sulfate, sodium chloride, and sodium bicarbonate purchased from Fisher Scientific. Reagents used for the extraction and analysis of carotenoids included HPLC grade ammonium acetate, ethyl acetate, glacial acetic acid, methanol, and water as well as ACS grade acetone, ethanol, hexanes, isopropanol, methyl tert-butyl ether, and petroleum ether purchased from Fisher Scientific. Authentic standards for α-carotene, β-carotene, lutein, lycopene, trans-β-apo-8′-carotenal, retinyl palmitate, and zeaxanthin were purchased from Sigma Aldrich.

### 1.2.2 Sorghum transformation, plant material, and harvest conditions

Immature embryo explants isolated from greenhouse grown sorghum plants were transformed with *Agrobacterium* auxotrophic strain LBA4404 Thy-carrying a ternary vector transformation system to generate transgenic sorghum plants as previously described (Che et al., 2018). Transgenic grains used in this study were generated from two transformation vectors, ABS4-1 and ABS4-2 (Table 1). ABS4-1 carried the maize codon-optimized phytoene desaturase *CRTI* (Ye et al., 2000; Che et al., 2016) gene from Pantoea ananatis to increase provitamin A biosynthesis, the maize codon-optimized phytoene synthase *PSY1* (Paine et al., 2005; Che et al., 2016) gene from *Zea mays* L. to modulate flux through the carotenoid pathway, the homogentisate geranylgeranyl transferase *HGGT* (Cahoon et al., 2003; Che et al., 2016) gene from *H. vulgare* to increase vitamin E accumulation, the maize codon optimized phytase *PhyA* (Han et al., 1999) gene from *Aspergillus niger* used for phytate metabolism, and the phosphomannose isomerase *PMI* (Negrotto et al., 2000; Che et al., 2018) gene from *Escherichia coli* as selectable marker. ABS4-2 carried all the identical genes as described in ABS4-1 except *PSY1* was replaced by maize codon-optimized *CRTB* (Neudert et al., 1998), another phytoene synthase gene from *Erwinia uredovora*. In both constructs, *CRTB*, fused to a maize codon optimized delta-4-palmitoyl-ACP desaturase gene transit peptide (Cs-DPAD) (Albertsen et al., 2014), and *PSY1* were driven with the same sorghum α-kafirin promoter (Abbitt et al., 2009). *CRTI* was fused to a maize codon optimized ribulose-1,5-bisphosphate carboxylase small subunit transit peptide (PS SSU TP) (Coruzzi et al., 1983; Albertsen et al., 2014) and was driven by the sorghum β-kafirin promoter (Abbitt et al., 2009). *PhyA* was fused to a *H. vulgare* alpha amylase signal peptide (BAASS) (Park et al., 2016) and was driven by the ZM-LEG1A promoter. HGGT was driven by the Zm-WS1 whole seed promoter (aka seed-specific KG86 promoter (Fu et al., 2010)), and *PMI* was driven with the Zm-UBI1 promoter (Negrotto et al., 2000; Che et al., 2018).

**Table 1.** Description of transgenic and non-transgenic material used in this study.

| Transgene<br>Combination<sup>z</sup> | Event<br>Identifier | Genetic<br>Background |
| --- | --- | --- |
| ABS4-1 = HGGT +<br>PSY1 +CRTI + PhyA | ABS4-1A<br>ABS4-1B<br>ABS4-1C | Tx430 |
| ABS4-2 = HGGT +<br>CRTB + CRTI + PhyA | ABS4-2A<br>ABS4-2B<br>ABS4-2C<br>ABS4-2D | Tx430 |
| NA | Control | Tx430 |

<sup>z</sup>Enzyme abbreviations are defined as follows: PSY1 phytoene synthase 1, CRTB bacterial phytoene synthase, CRTI bacterial phytoene desaturase, HGGT homogentisate geranylgeranyl transferase, PhyA phytase, and PSY1 phytoene synthase 1

Transgenic events derived from the same transformation vector as shown in Table 1 represent independent transgenic events. All transgenic events used in this study were homozygous with a single T-DNA insertion. Both null segregants isolated through segregation of corresponding transgenic events and wild-type Tx430 were used as controls. Panicles were collected from sorghum plants 45-day after pollination, air dried at 24 °C for 2 weeks before threshing, and then stored at −80 °C.

### 1.2.3 High-Throughput Sorghum Flour and Experimental Porridge Preparation

Whole sorghum kernels were added to fill approximately half of a 15 mL polycarbonate vial (SPEX Sample Prep, Metuchen, NJ, USA; UX-04545-71) containing two ¼” 440C stainless steel balls (Grainger, Minooka, IL, USA; 4RJJ1). Samples were milled into flour using a GenoGrinder 2010-115 (SPEX Sample Prep) operated at 1400 RPM for 5 minutes. Sorghum flour was stored at -40° C until analysis.

Steeped samples underwent the following pre-treatment: Sorghum flour (0.4 g) was suspended in 0.8 mL of doubly distilled water and incubated at 37° C for 3 hours at 120 oscillations per minute (OPM). After incubation, an additional 0.8 mL of doubly distilled water was added to each tube and samples were agitated by hand. Samples were then cooked and processed as described below.

Porridges were formulated based on a traditional Burkina-Faso style toL comprised of 20% sorghum flour and 80% water by weight (Da et al., 1981) and scaled to a 10 mL volume to align with the cooking method outlined by Lipkie and others (Lipkie et al., 2013). Briefly, 0.4 g of sorghum flour was weighed into 15 mL flip-cap tubes and 1.6 mL of doubly distilled water was added. Each tube was agitated by hand until flour was fully dispersed. Tubes were transferred to metal racks with their lids open and racks were placed into an induction heated stock pot submerging only the portion of the tubes that contained porridge in water. The stockpot lid was replaced to maintain temperature at 100° C and high humidity while the samples cooked for 10 minutes. After cooking, racks were removed and 100 mg (± 5 mg) of canola oil was added to each tube (5% lipid by porridge mass) and incorporated by manual mixing. After the addition of canola oil (∼30 min), sample tubes were transferred to a -80° C freezer and subjected to a simulated digestion within 12 hours. All sorghum samples were processed in triplicate to account for experimental variation.

### 1.2.4 High-Throughput *in vitro* Digestions of Sorghum Porridge

A previously described three-stage high-throughput *in vitro* digestion method that utilizes a Tecan Freedom EVO 150 liquid handling system (Tecan; Mannedorf, Switzerland) was adapted for sorghum porridge (Lipkie et al., 2013; Hayes et al., 2020). Briefly, thawed porridge samples were mixed with an oral phase solution (containing 31.8 mg/mL of ⍰-amylase) and incubated at 37° C for 10 minutes at 120 OPM. Samples were diluted with a 10 mg/mL pepsin solution (in 0.1 M HCl) and saline (0.9% NaCl) solution and then adjusted to pH 2.5 by addition of 1.0 M HCl. Sample tubes were blanketed with nitrogen gas and incubated at 37° C for 1 hour at 120 OPM. Samples were adjusted to pH 5.0 using 1.0 M NaHCO_3_. Then, pancreatin-lipase (20 mg/mL of each enzyme) and bile extract (30 mg/mL) solutions were added to each sample. Samples were adjusted to pH 7.0 using 1.0 M NaHCO_3_ and saline was added to dilute samples to a final volume of 10 mL. Nitrogen gas was added to each tube and samples were incubated at 37° C for 2 hours at 120 OPM. After incubation, samples were hand-agitated and 4 mL of digesta was removed, aliquoted, and stored at -80° C for future analysis. The remaining samples were blanketed with nitrogen gas and centrifuged at 3,428 *g* for 75 minutes in an Eppendorf 5920R centrifuge (Eppendorf, Hamburg, Germany) maintained at 4° C. Following centrifugation, 4 mL aliquots of the aqueous fraction were filtered through 0.22 µm cellulose acetate filters. Samples were blanketed with nitrogen gas and stored at -80° C until analysis.

### 1.2.5 High-Throughput Extraction and Analysis of Carotenoids

#### 1.2.5.1 Extraction of Carotenoids from Sorghum Flour

A rapid extraction protocol was developed based on a previously validated method for extracting carotenoids from sorghum flour (Lipkie et al., 2013). Briefly, 300 mg of sorghum flour was weighed into 2.0 mL microfuge tubes. Two 1/8” 440C stainless steel balls (Grainger; 4RJH5) were added and 150 µL of HPLC grade water was added into each tube to hydrate the sample matrix. After resting for 5 minutes, 30 µL of 150 µM retinyl palmitate dissolved in ethanol was spiked into each sample as an internal standard. Samples rested for an additional 10 minutes prior to the addition of 1 mL of 3:2 acetone:ethyl acetate + 0.01% BHT (*w*/*v*). Tubes were capped, placed in sample racks, and shaken at 1400 RPM for 45 seconds in a GenoGrinder 2010-115. Samples were centrifuged at 20,000 *g* for 3 minutes and the supernatant was collected into borosilicate culture tubes. The remaining sample pellets were extracted as outlined above two more times, but with methyl tert-butyl ether + 0.01% BHT (*w*/*v*) substituted for the final extraction solvent to better capture non-polar carotenoids such as all-*trans*-β-carotene. The combined supernatants were dried using a RapidVap (Labconco, Kansas City, MO, USA), blanketed with nitrogen, and stored at -80° C. Samples were redissolved in 2 mL of 1:1 methanol:ethyl acetate and filtered through 0.45 µm PTFE syringe filters prior to analysis. All sorghum flour samples were extracted in triplicate and estimates represent an average. Extraction procedures were conducted under low-light to minimize carotenoid photoisomerization.

#### 1.2.5.2 Extraction of Carotenoids from Digesta and Aqueous fractions

Extraction from digesta and aqueous fractions was based on a previously validated method and adapted for a Tecan Freedom EVO 150 liquid handling unit (Lipkie et al., 2013; Hayes et al., 2020). Briefly, 100 µL of retinyl palmitate dissolved in ethanol was added to digesta (150 µM retinyl palmitate) and aqueous samples (30 µM retinyl palmitate) containing 2 or 4 mL of starting material, respectively. Samples were extracted three times with one volume of 1:3 acetone:petroleum ether + 0.01% BHT (*w*/*v*). Between extractions, samples were vortexed for 1 minute, centrifuged for 2 minutes at 4,000 RPM, and the supernatant was removed and saved. Combined supernatants were dried in a RapidVap and stored at -80° C until analysis (<24 hours later). Digesta and aqueous fraction samples were resolubilized in 200 or 100 µL of 1:3 ethyl acetate:methanol + 0.01% BHT (*w*/*v*), respectively, and filtered with a 0.22 µm PTFE filter prior to analysis.

#### 1.2.5.3 Analysis of Carotenoids

Carotenoids were analyzed using a modified high performance liquid chromatography photo diode array detector (HPLC-PDA) method developed by Lipkie and colleagues (Lipkie etmal., 2013). Briefly, filtered carotenoid extracts were run on a Waters Alliance e2695 (Waters Corporation, Milford, MA, USA) with mobile phases containing HPLC grade 98:2 methanol: 2% 1.0 M ammonium acetate adjusted to pH 4.6 (A) and ethyl acetate (B). Solvent flow rate was maintained at 0.45 mL/minute and carotenoids were separated on a 2.0 mm x 150 mm YMC C_30_ column with 3 µm particle size (YMC America, Devens, MA) using the following gradient: 100% A to 20% A over 4.93 minutes, 20% A to 0% A over 1.65 minutes, 0% A held for 1.46 minutes, a return to 100% A over 1.0 minute, and a hold on 100% A for 3.46 minutes to recondition the column. The column compartment and autosampler were maintained at 35 and 15° C, respectively. Carotenoids were quantified at 450 nm using a Waters 2998 PDA detector based on response curves of authentic standards. For analytes quantified without authentic standards, an adjusted slope was calculated using a ratio of molar extinction coefficients with the most structurally related carotenoid available. The internal standard retinyl palmitate was quantified at 325 nm.

### 1.2.6 Extraction and Analysis of Minerals from Sorghum Flour

Minerals were extracted from 10 g of sorghum flour using 20 mL of a diethylenetriaminepentaacetic acid (DTPA) solution containing 5 mM DTPA, 0.1 M triethanolamine, and 10 mM calcium chloride dihydrate. Samples were shaken at room temperature at 160 OPM for 2 hours. After shaking, samples were immediately filtered through Ahlstrom 624 filter paper into 15 mL tubes and analyzed using ICP-OES. All extractions and analyses were conducted by A&L Great Lakes Laboratories (Fort Wayne, IN, USA).

### 1.2.7 Extraction and Analysis of Inositol Phosphates

Extraction of inositol phosphates from raw and steeped sorghum samples was carried as described previously with minor modifications (Zhang et al., 2017). Briefly, 50 mg of ground sorghum samples were defatted using 1.5 mL of hexane. The hexane layer was carefully removed after centrifuging (3600 rpm, 4 min) and the pellet was dried under nitrogen. The dried samples were then resuspended with 1 mL of HCl (0.4 M) and sonicated for 1 h at room temperature. Samples were centrifuged, supernatants were transferred into culture tubes and dried using RapidVap vacuum evaporator. Dried samples were then resolubilized with 500 ul of 5% acetonitrile, filtered using 0.45-µm injected µm PES syringe filter and injected for LC-MS analysis.

Inositol phosphate content was determined using a Waters UPLC Acquity I Class system coupled with a Xevo TQ-S triple quadrupole mass spectrometer. Separation was performed on a BEH C18 column (1.7 µm, 2.1 mm x 50 mm) at a flow rate of 0.5 mL/min. Mobile phases consisted of (A) 5% aqueous acetonitrile containing ion pairing agents (5mM dihexylammonium acetate and 5 mM ammonium acetate) and (B) 100% acetonitrile. The elution gradient was adapted and optimized for new column particle size, geometry, and system dead volume (Zhang et al., 2017). Inositol phosphates were identified by comparing retention time and molecular mass of peaks with those of authentic standards. Calibration curves of D-myo-Inositol-1-monophosphate dipotassium salt (IP1), D-myo-Inositol-1,4-diphosphate sodium salt (IP2), D-myo-Inositol-1,4,5-triphosphate, sodium salt (IP3), D-myo-Inositol-1,3,4,5-tetraphosphate, sodium salt (IP4), D-myo-Inositol-1,3,4,5,6-pentaphosphate, sodium salt (IP5) and myo-Inositol-hexakis (dihydrogen phosphate) (IP6) were used to quantify inositol phosphate species using optimized multiple reaction monitoring experiments. Analytical conditions were maintained as follows: source: electrospray ionization in negative mode; capillary voltage: 3.0 kV; probe temp: 150 °C; source temp: 600°C; desolvation gas flow: 1000 L/hr; cone gas flow: 5 L/hr.

### 1.2.8 Statistical Analysis and Data Visualization

Statistical analyses and data visualization were conducted using R version 4.1.1 (R Development Core Team, 2018). Box and whisker plots were generated using ggplot2 using the ‘Wes Anderson’ color palette generator (Ram et al., 2018; Wickham, 2016). Fixed-effect analysis of variance models were generated to determine if genetic background, transgene state of the transgenic construct, or phytase activation pre-treatment significantly affected outcomes measured in our experiments. For the analysis of raw material carotenoid, mineral, and phytate content, (excluding pre-treated phytase activated samples) the following model was used:

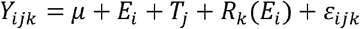

*Y*_*ijk*_ represents the estimate for a given analyte within *i*^th^ event, *j*^th^ transgene state, and *k*^th^ technical replicate. µ represents the mean of a given analyte or metric, *E*_*i*_ represents the contribution due to genetic factors for *i*^th^ event of the germplasm, T represents the contribution due to the transgene state of the transgenic construct for the *j*^th^ allele, *R*_*k*_ (*E*_*i*_) represents the contribution of the *k*^th^ technical replicate within transgenic event, and can be interpreted as within event variation, and *ε*_*ijk*_ represents the residual error. If significance was determined, a Tukey-Kramer post-hoc test (α = 0.05) was conducted to determine between which groups differences exist using the package agricolae (Mendiburu & Yanseen, 2021).

To isolate the effect of transgenic construct transgene state or phytase activation pre-treatment on relative and absolute bioaccessibility, the following simplified models were used:

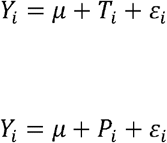

Where model parameters are as previously defined and *P*_*i*_ indicates phytase activation pre-treatment. Tukey-Kramer post-hoc tests (α = 0.05) were used to define significance between groups within a given analyte.

To determine significance among the interactions between transgenic construct transgene state and phytase activation pre-treatment, the following model was used:

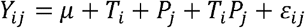

Where model parameters are as previously defined and *T*_*i*_ *P*_*j*_ indicates the interaction term between transgene state and phytase activation pre-treatment.

## 1.3 Results

### 1.3.1 Carotenoid content increased in transgenic sorghum events

Sorghum flours were analyzed for several major carotenoids including those with provitamin A activity (Table 2). A comprehensive summary and statistical analysis of all carotenoids and their geometric isomers is detailed in Table S1. Carotenoid values are presented on a “per serving” basis to reflect the amount present in a standard 200 g serving of porridge. Additionally, we report multiple aggregate values including xanthophylls (lutein, zeaxanthin, 11-cryptoxanthin, and β-cryptoxanthin), carotenes (⍰-carotene, *cis*-β-carotene, all-*trans*-β-carotene, and all-*trans*-lycopene), and provitamin A content (1/2(⍰-cryptoxanthin + β-cryptoxanthin + ⍰-carotene + *cis*-β-carotene) + all-*trans*-β-carotene). Overwhelmingly, transgenic material outperformed null segregants and the wild-type control in terms of carotenoid content. Depending on the analyte, transgenic events contained on average 266% to 3215% more carotenoids than their non-transgenic counterparts and the wild-type control. A 200 g porridge serving made with the null segregants or wild type control used in this study contained on average 0.15 mg of xanthophylls, 0.03 mg of carotenes, and 0.02 mg of provitamin A content while a serving of porridge made with transgenic events contained on average 0.35 mg of xanthophylls, 0.70 mg of carotenes, and 0.67 mg of provitamin A content. By design, transgenic sorghum events were particularly high in all-*trans*-β-carotene with an average content of 0.63 mg per 200 g serving of porridge.

**Table 2:**
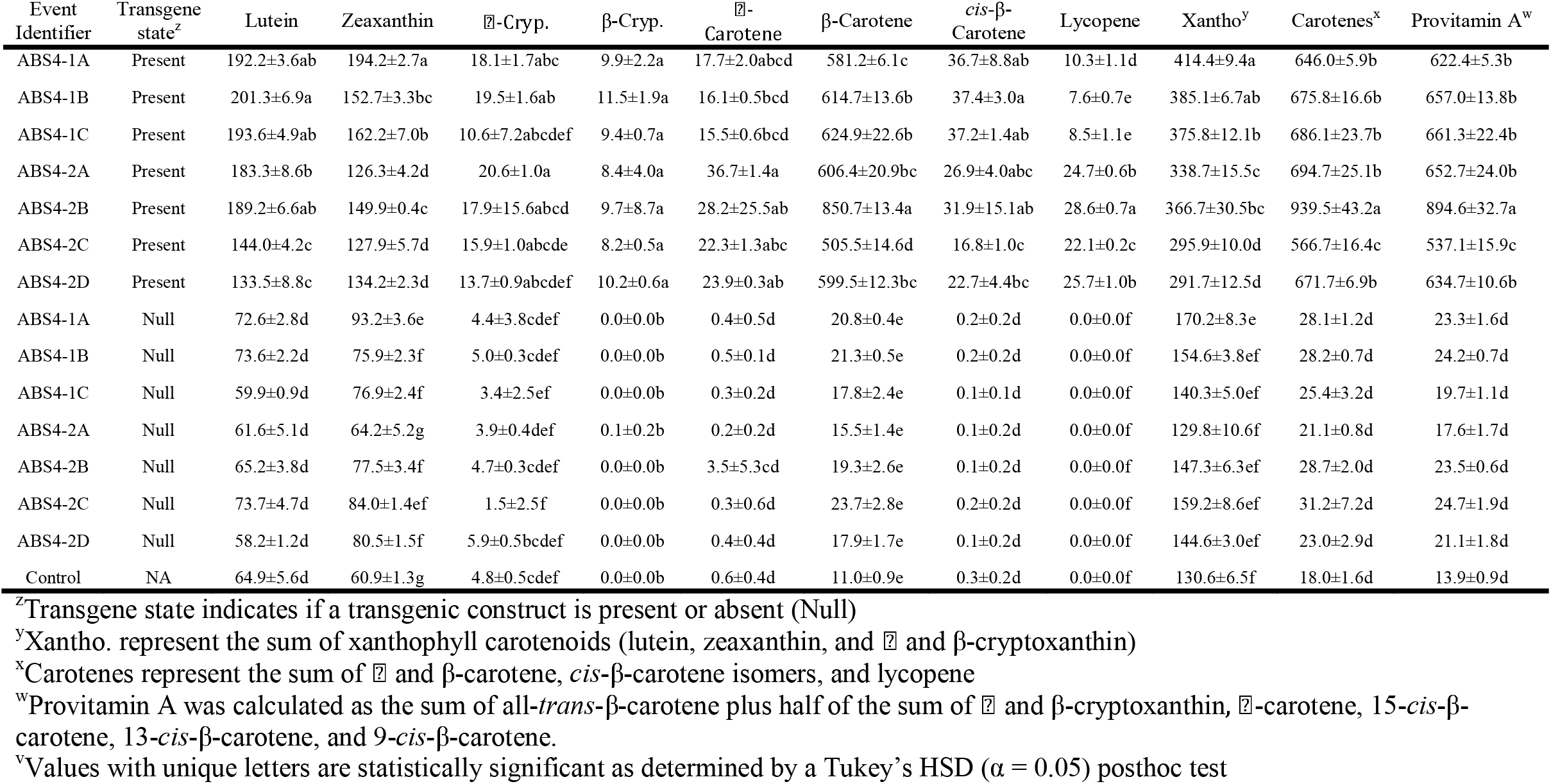
Carotenoid profiles of raw material reported as means ± standard deviation (µg/200 g porridge (40 g dry flour))

### 1.3.2 Relative and absolute bioaccessibility of carotenoids varied in transgenic and non-transgenic sorghum events

Relative bioaccessibility is defined as the proportion of a given carotenoid incorporated into absorbable bile salt-lipid micelles compared to the amount present in the digesta fraction. In our germplasm, relative bioaccessibility was higher for almost all carotenoids in the non-transgenic material (Fig. 1). On average, 23.5% of all-trans-β-carotene in null segregants and wild type control was bioaccessible compared to 18.8% in transgenic lines. A detailed breakdown of relative bioaccessibility values for all carotenoids and respective geometric isomers quantified are reported in Table S2. β-Cryptoxanthin, ⍰-carotene, and all-*trans*-lycopene were low or not detectable in non-transgenic material (Fig. 1; Table 2). No significant differences in relative bioaccessibility for all-*trans*-β-carotene were seen amongst the non-transgenic material while some transgenic lines (e.g. ABS4-1C) exhibited values statistically similar to their non-transgenic counterparts (23.6%; Table S2).

Absolute bioaccessibility is defined as the quantity of absorbable carotenoids released during digestion and normalized to the amount present in the starting material. For all carotenoids measured in our study, transgenic material had substantially higher absolute bioaccessibility (Fig. 2). On average, transgenic events released 0.⍰8 mg of all-*trans*-β-carotene while non-transgenic material released 4.57 µg in a 200 g serving of porridge. This finding represents a 2582% increase in absorbable all-*trans*-β-carotene from porridges made using transgenic sorghum events. Individual transgenic and null segregants exhibited differences in bioaccessibility within their respective groups. Notably, ABS4-2A released the most all-*trans*-β-carotene, total xanthophylls, carotenes, and provitamin A carotenoids (Table S2).

**Figure 1.**
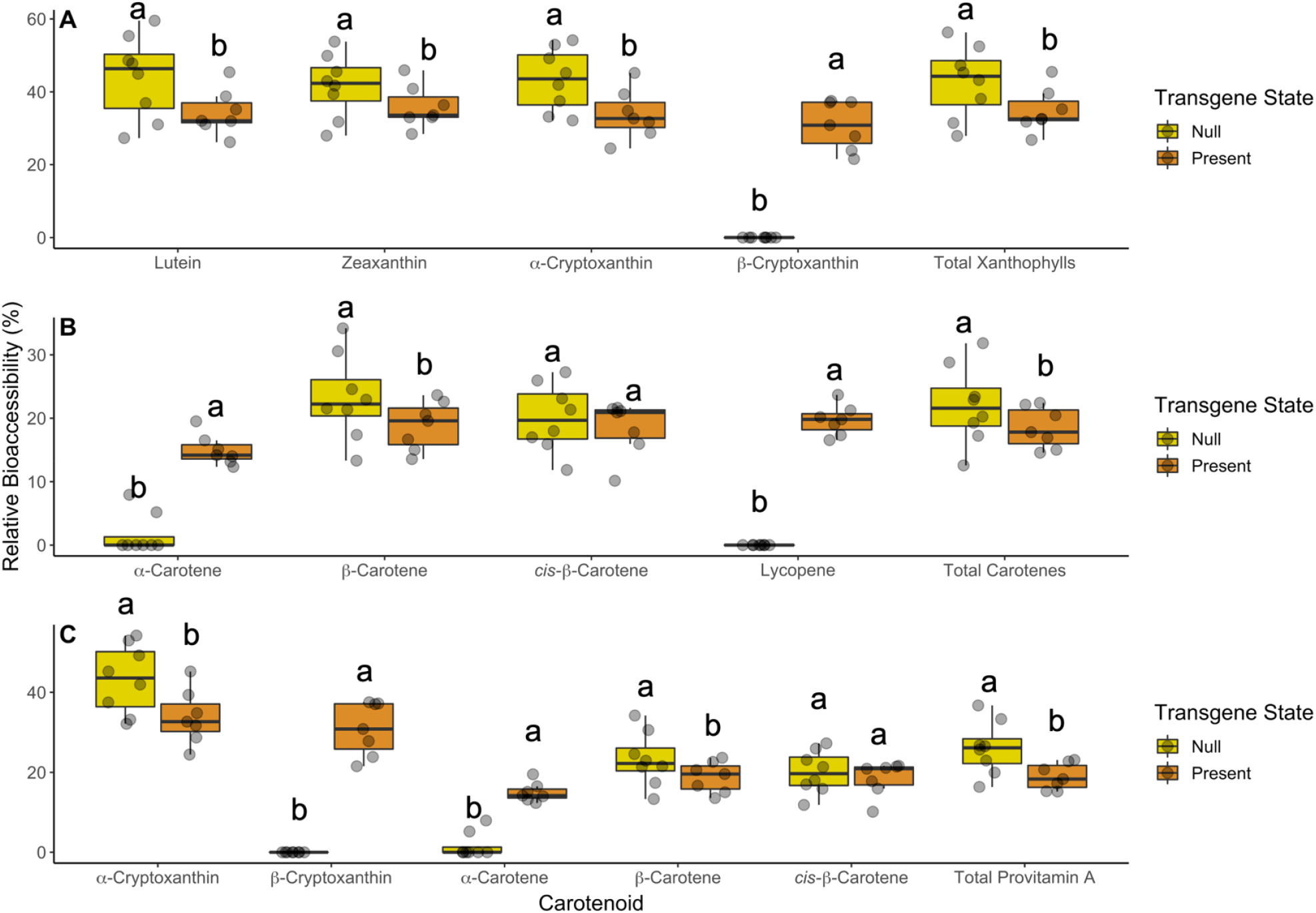
Relative bioaccessibility comparing transgenic to non-transgenic material. Plots are separated by (A) xanthophylls, (B) carotenes, and (C) provitamin A carotenoids. Lowercase letters above box and whisker plots that are different indicate statistical significance within a carotenoid as determined by a post-hoc Tukey’s HSD test. Transgene state indicates if a transgenic construct is present or absent (Null).

**Figure 2.**
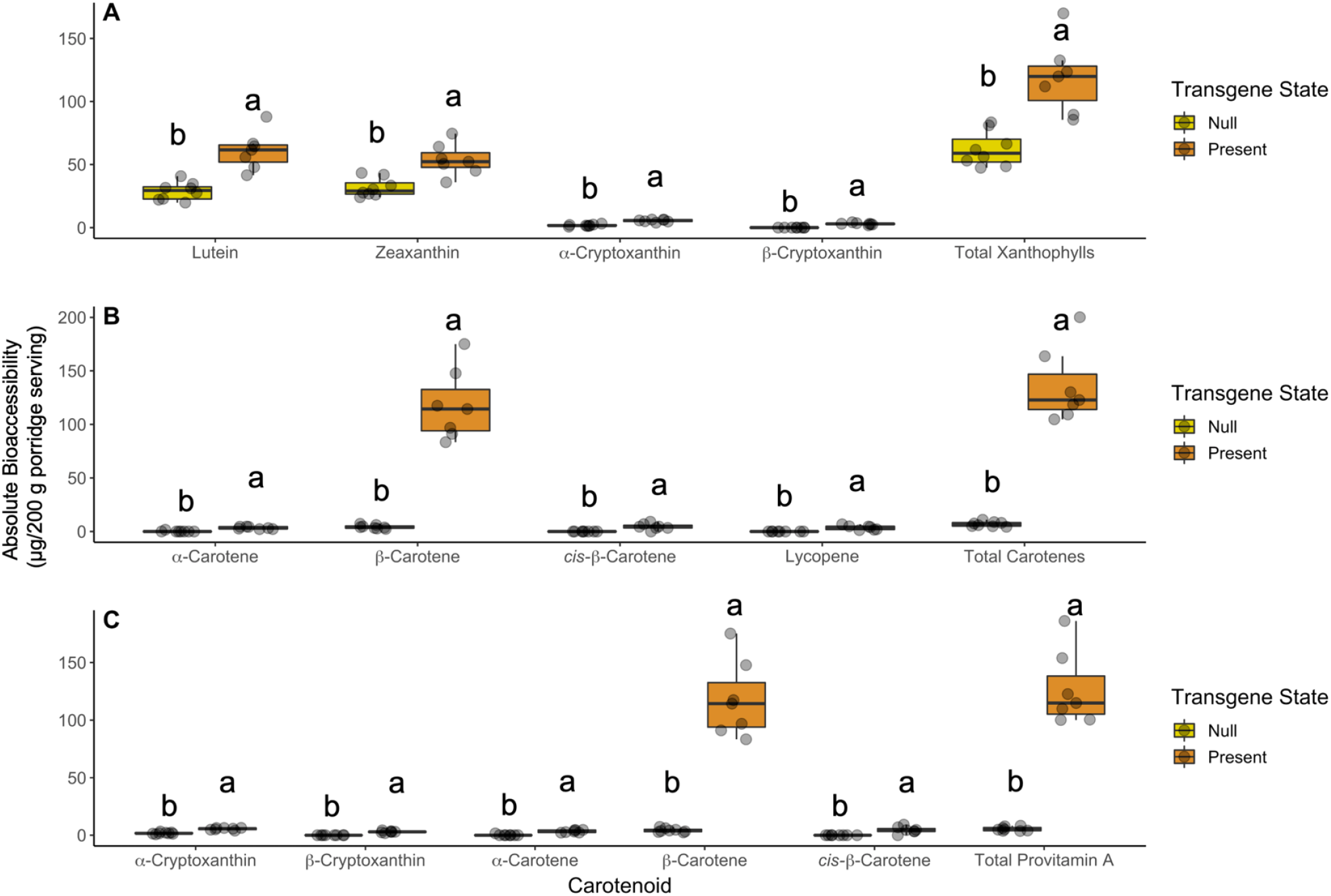
Absolute bioaccessibility comparing transgenic to non-transgenic material. Plots are separated by (A) xanthophylls, (B) carotenes, and (C) provitamin A carotenoids. Lowercase letters above box and whisker plots that are different indicate statistical significance within a carotenoid as determined by a post-hoc Tukey’s HSD test. Transgene state indicates if a transgenic construct is present or absent (Null).

### 1.3.3 Mineral profiles were similar in all sorghum lines studied

Mineral elements common to cereal grains were analyzed in sorghum samples to better understand the potential interactions between their presence and carotenoid bioaccessibility and to determine if specific events caused alterations in mineral homeostasis (Table 3, Table S3). Among these minerals were divalent cations essential for human health and well-being such as iron, zinc, and calcium. Mineral profiles of all sorghum lines did not significantly vary regardless of transgene state. It is important to emphasize that these constructs were not engineered to differentially accumulate minerals and all germplasm studied here were grown in a controlled environment.

**Table 3:** Mineral concentrations in sorghum flour reported as means ± standard deviation (mg/200 g porridge (40 g dry flour))

| Event Identifier | Transgene state<sup>z</sup> | Zinc | Iron | Calcium | Magnesium |
| --- | --- | --- | --- | --- | --- |
| ABS4-1A | Present | 1.3±0.0abc<sup>y</sup> | 1.4±0.2b | 14.7±2.3ab | 58.7±6.1abc |
| ABS4-1B | Present | 1.6±0.3abc | 1.5±0.1b | 16.0±0.0ab | 68.0±13.9ab |
| ABS4-1C | Present | 1.5±0.2abc | 1.8±0.6ab | 17.3±2.3ab | 65.3±6.1ab |
| ABS4-2A | Present | 2.0±0.2a | 2.1±0.7ab | 17.3±6.1ab | 73.3±4.6a |
| ABS4-2B | Present | 1.6±0.4ab | 1.3±0.1b | 16.0±0.0ab | 68.0±0.0ab |
| ABS4-2C | Present | 1.6±0.2abc | 1.7±0.3ab | 16.0±0.0ab | 66.7±12.9ab |
| ABS4-2D | Present | 1.5±0.1abc | 1.6±0.2ab | 17.3±2.3ab | 70.7±4.6ab |
| ABS4-1A | Null | 1.4±0.1abc | 1.3±0.1b | 13.3±2.3b | 61.3±2.3abc |
| ABS4-1B | Null | 1.3±0.2abc | 1.5±0.2b | 21.3±6.1ab | 62.7±11.5ab |
| ABS4-1C | Null | 1.3±0.2abc | 1.4±0.5b | 13.3±2.3b | 57.3±6.1abc |
| ABS4-2A | Null | 1.5±0.2abc | 2.5±0.7a | 22.7±2.3a | 68.0±4.0ab |
| ABS4-2B | Null | 1.4±0.1abc | 1.5±0.1b | 16.0±0.0ab | 65.3±2.3ab |
| ABS4-2C | Null | 1.2±0.1bc | 1.4±0.1b | 13.3±2.3b | 56.0±8.0bc |
| ABS4-2D | Null | 1.8±0.2ab | 2.0±0.2ab | 17.3±2.3ab | 64.0±8.0ab |
| Control | NA | 1.0±0.3c | 1.3±0.2b | 13.3±2.3b | 45.3±2.3c |

<sup>z</sup>Transgene state indicates if a transgenic construct is present or absent (Null)
<sup>y</sup>Values with unique letters are statistically significant as determined by a Tukey's HSD ( $\alpha =$ 0.05) posthoc test

### 1.3.4 Total inositol phosphate pools decreased as phytate (IP6) was metabolized into other inositol phosphate forms (IP1-IP5) in steeped transgenic sorghum events

In addition to profiling minerals in the sorghum germplasm evaluated here, we also determined the effects of activating both native and heterologous phytase enzymes on inositol phosphate (IP1-IP6) profiles and carotenoid bioaccessibility. Without steeping prior to porridge production and digestion, transgenic and non-transgenic events did not substantially differ from one another in terms of their phytate metabolite (IP1-IP5) profiles (Table S4). However, large differences were observed as a function of steeping (Fig. 3, Table S4). Phytate metabolites were significantly higher in steeped transgenic material compared to their non-steeped counterparts as well as both steeped and non-steeped null segregants and non-transgenic controls. In non-steeped transgenic material, IP1-IP3 represented 0.6% of the total inositol phosphate pool on average whereas IP1-IP3 comprised 25.7% of the total inositol phosphate pool in steeped transgenic events. In non-transgenic material, changes in inositol phosphate profiles were less pronounced, but statistical differences were detected as a function of steeping in IP2-IP6 (Table S4). Additionally, an overall reduction in total inositol phosphate species was observed in transgenic material after steeping compared to non-steeped counterparts as well as both steeped and non-steeped null segregants and wild-type control (Fig. 3, Table S4). More precisely, steeped transgenic material contained, on average, 370.8 mg of total inositol phosphates whereas non-steeped events contained 937.1 mg per 200 g of porridge.

**Figure 3.**
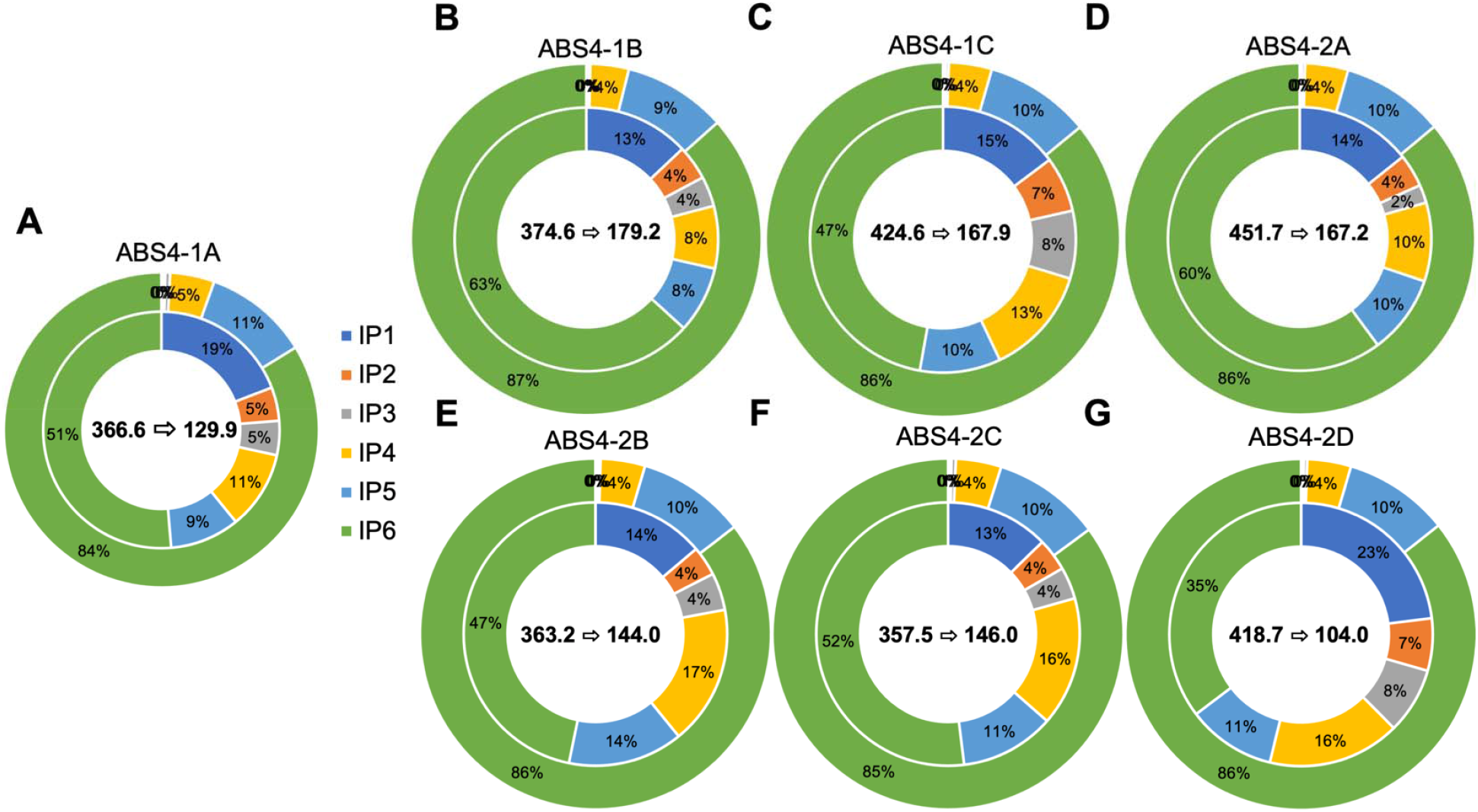
Donut plots of phytate (IP6) and its metabolites (IP1-IP5) in transgenic events represented as a percentage of total inositol phosphates (IP1-IP6). The outer rings represent non-steeped porridges while the inner rings represent steeped porridges. The average change in total inositol phosphates as a function of steeping for each event are displayed in the center of its corresponding figure. The first number represents total inositol phosphates in the non-steeped porridges while the latter number represents total inositol phosphates after steeping mg/200 g porridge (40 g dry flour)).

### 1.3.5 The molar ratios of phytate to iron and zinc were reduced in steeped transgenic sorghum events

We calculated molar ratios of phytate (IP6) to zinc and iron as a substitute for mineral bioaccessibility. In this analysis, both steeped and non-steeped transgenic and non-transgenic material were compared to determine phytate degradation efficiency among wild-type, transgenic and corresponding nulls. As shown in Fig. 4 and Table S5, no significant differences in the phytate to zinc (21.6 and 21.2; transgenic and non-transgenic) and phytate to iron (18.2 and 16.4; transgenic and non-transgenic) ratios were observed for all the materials tested without steeping. However, a significant reduction in phytate to zinc (4.9 and 17.6; transgenic and non-transgenic) and phytate to iron (4.1 and 13.6; transgenic and non-transgenic) ratios was observed for all the sorghum lines after steeping, indicating that the steeping treatment was able to activate not only the heterologous phytase, but also naturally occurring phytase enzymes. Although steeping reduced the phytate to iron and zinc ratios in both transgenic and non-transgenic material, transgenic material exhibited the most dramatic reduction falling below 5.0 for both minerals (Fig. 4 and Table S5).

**Figure 4.**
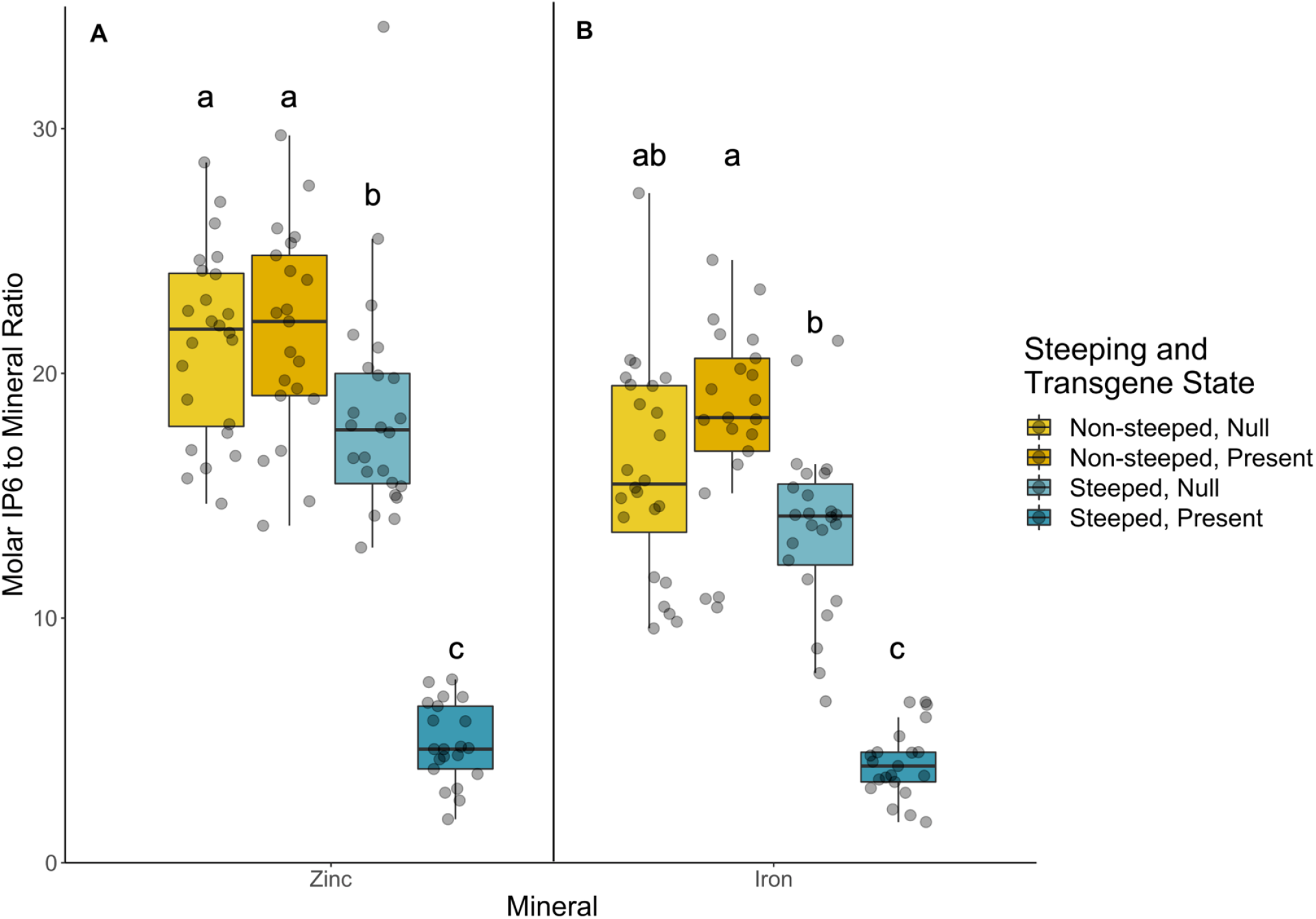
Molar IP6 to mineral ratios of transgenic and non-transgenic sorghum as well as activated or non-activated pre-treatments. The plot is separated by mineral class (A: zinc and B: iron) and lowercase letters above box and whisker plots that are different indicate statistical significance within a mineral as determined by a post-hoc Tukey’s HSD test. Phytase activation indicates if sorghum flours were pre-steeped prior to porridge production. Transgene state indicates if a transgenic construct is present or absent (Null).

### 1.3.6 Relative and absolute bioaccessibility of carotenoids were not affected by phytase activation in transgenic sorghum events

As divalent minerals have been shown to impact carotenoid bioavailability (Corte-Real et al., 2016, 2017, 2018), it was critical to determine if carotenoid bioaccessibility was affected by phytase activation wherein divalent cations normally bound to phytate would be liberated and interfere with micelle formation. For this comparison, we focused our analysis on the transgenic events given the notable differences in their total phytate, likely due to the heterologous phytase activated by steeping (Fig. 3, Table S4). Relative bioaccessibility was not significantly impacted for any of the carotenoids measured in this study by phytase activation (Fig. 5). In terms of absolute bioaccessibility, *cis*-β-carotene isomers were statistically more bioaccessible from non-steeped wild-type and null controls, but no other individual or aggregate measurements of carotenoids were affected (Fig. 6, Table S2). A modest and consistent pattern can be seen in Fig. 6 and Table S2 suggesting a slight downward trend in deliverable carotenoids as a function of steeping and phytase activation, but these differences did not reach a level of statistical significance. Therefore, delivery of carotenoids and provitamin A carotenoids were found to be similar between steeped and non-steeped transgenic material and not significantly impacted by the presumed increase of divalent cations liberated through phytase activation.

**Figure 5.**
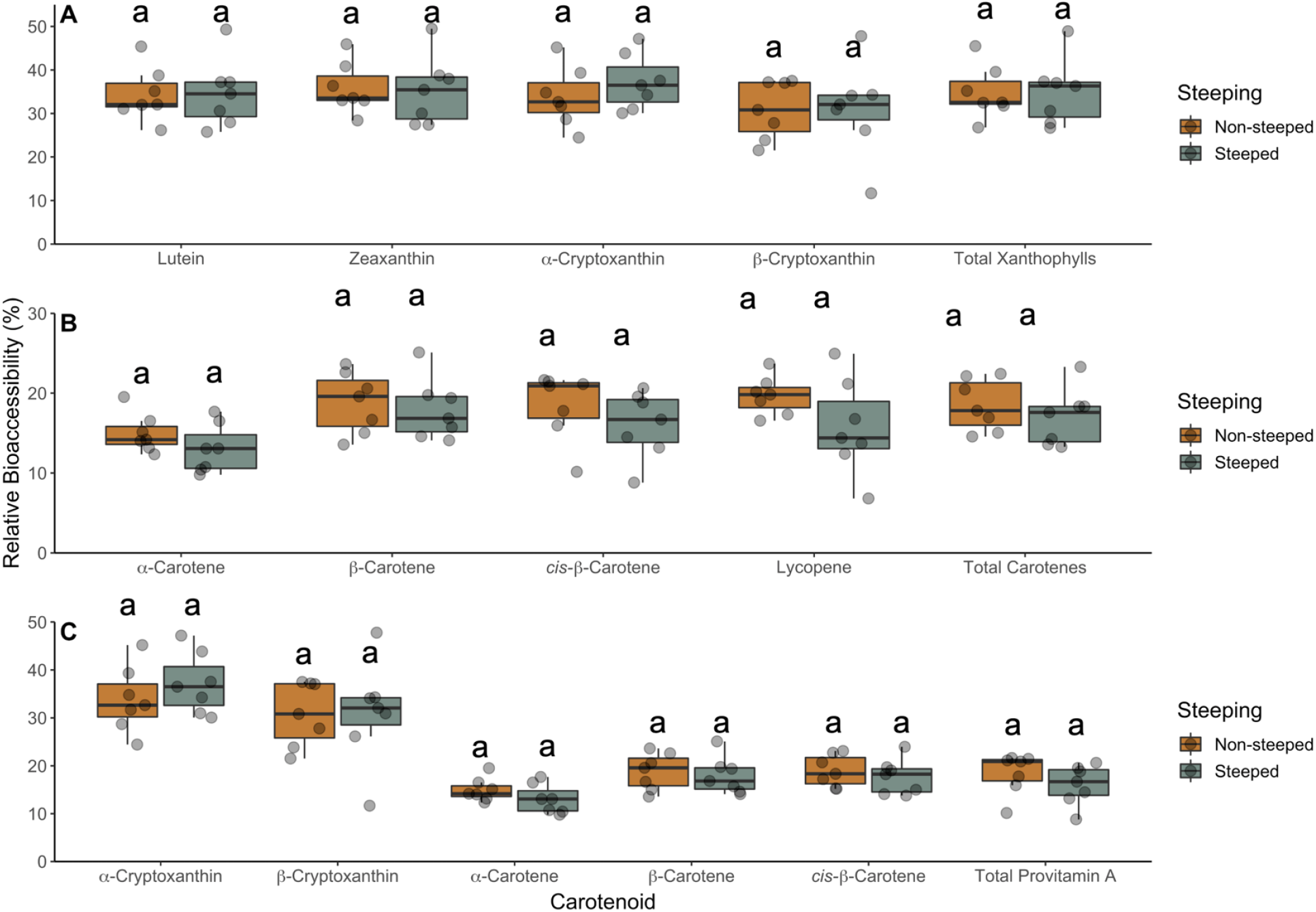
Relative bioaccessibility comparing phytase activated to non-activated material. Plots are separated by (A) xanthophylls, (B) carotenes, and (C) provitamin A carotenoids. Lowercase letters above box and whisker plots that are different indicate statistical significance within a carotenoid as determined by a post-hoc Tukey’s HSD test. Phytase activation indicates if sorghum flours were pre-steeped prior to porridge production.

**Figure 6.**
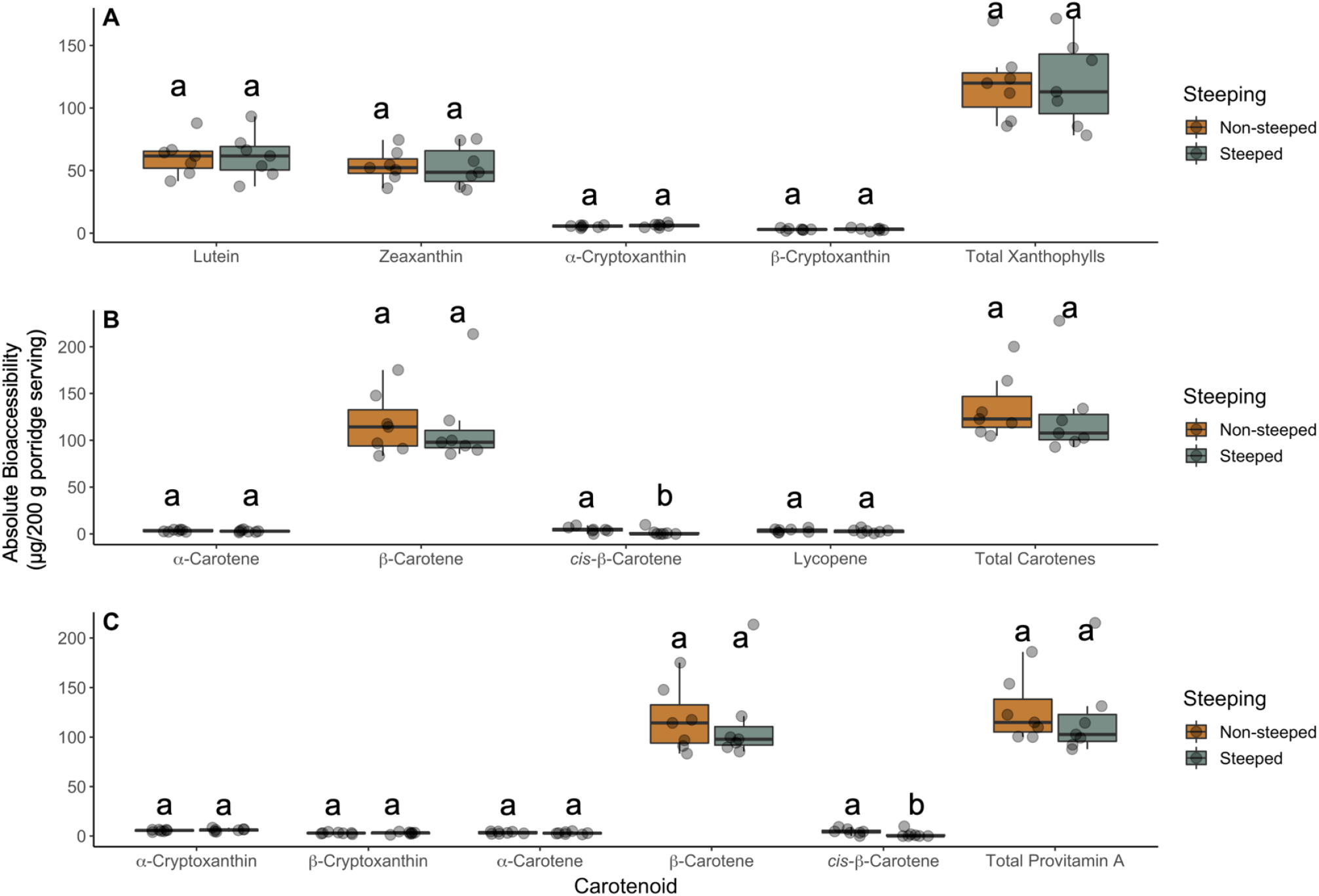
Absolute bioaccessibility comparing phytase activated to non-activated material. Plots are separated by (A) xanthophylls, (B) carotenes, and (C) provitamin A carotenoids. Lowercase letters above box and whisker plots that are different indicate statistical significance within a carotenoid as determined by a post-hoc Tukey’s HSD test. Phytase activation indicates if sorghum flours were pre-steeped prior to porridge production.

## 1.4 Discussion

Among the leading causes of death for children under 5 years of age in sub-Saharan Africa are vitamin A and mineral deficiencies (Black et al., 2008). The sorghum germplasm used in this study is part of long-term efforts to address nutritional deficiencies in the developing world. Events were specifically engineered to address how modifying the carotenoid biosynthetic pathway in endosperm tissue impacts carotenoid profiles and bioaccessibility. Secondarily, the interactions among carotenoid bioaccessibility, phytase activation, and the presence of liberated divalent cations was also studied by including PhyA in these constructs. The two constructs used in this study differed only in that ABS4-1 contained PSY1 while ABS4-2 contained CRTB (Table 1). In both cases, carotenoid profiles in the germplasm represent improvements over previous iterations with prominent increases in all-*trans*-β-carotene (Lipkie et al., 2013; Che et al., 2016, 2019). However, concentrations of xanthophylls such as lutein and zeaxanthin were slightly lower in this germplasm likely due to increased pathway flux towards all-*trans*-β-carotene. Given that PSY1 and CRTB both encode phytoene synthase and catalyze the same reaction through conserved prenyl transferase domains, changes in gene expression through codon optimization (codon optimized PSY1 was used in this study compared to previous iterations (Che et al., 2016)) and/or difference in the enzyme catalytic activity differences may have contributed to variation in xanthophyll content as well as provitamin A carotenoids in different transgenic constructs (Cunningham & Gantt, 1998). Regardless, xanthophyll concentrations in transgenic events were between ∼3-4x higher than our non-transgenic control (Tx430) and may provide greater long term benefits related to eye health (Table 2) (Mares, 2016; Thomas & Johnson, 2018).

Depending on factors such as age and sex, the estimated average requirement (EAR) of retinol activity equivalents (RAE) for vitamin A can range from 275 µg up to 885 µg for children 4-8 and lactating mothers, respectively (Institute of Medicine (US) Committee on Nutrition Standards for National School Lunch and Breakfast Programs, 2008). The reported conversion of dietary all-*trans*-β-carotene is 12:1 (Trumbo et al., 2001). However, biofortified cereals like maize and Golden rice have been shown to have more effective conversion rates (6:1 and 3.8:1, respectively) (Tang et al., 2009; Li et al., 2010). Given the similarity of sorghum flour’s composition to maize, retinol conversion rates may be closer to those reported for other cereals. A recent report using a Mongolian gerbil model suggests that the conversion efficiency of all-*trans*-β-carotene derived from similar transgenic sorghum events to retinol is 4.5:1 (You, 2016). Based on the 4.5:1 conversion rate, a 200 g porridge serving made with the transgenic events used in this study contains on average 665.69 µg of β-carotene equivalents; fulfilling 53.7% of a 4–8-year-old child’s vitamin A EAR (275 µg RAE). However, the efficiency of provitamin A carotenoid conversion to retinol will be affected by many other factors, such as porridge matrix, preparation method, carotenoid bioaccessibility, and interindividual differences in absorption and metabolism capabilities (Weber & Grune, 2012).

Consistent with previous reports, relative bioaccessibility (micellarization efficiency) was lower in transgenic events with high concentrations of carotenoids (Fig. 1) (Failla et al., 2012; Lipkie et al., 2013; Aragón et al., 2018). This trend could not be observed in β-cryptoxanthin, ⍰-carotene, and lycopene as these carotenoids were low or not detectable in non-transgenic material. We hypothesize that due to the prominently higher carotenoid concentrations in transgenic events, the process by which carotenoids partition into water-soluble micelles may have been over-saturated considering the limiting factor of lipid content in these models.

Regardless, the transgenic events used in this study have the potential to deliver significantly more total and provitamin A carotenoids than their non-transgenic counterparts (Fig. 2), with a particular emphasis on all-*trans*-β-carotene. Despite the modestly lower relative bioaccessibility in these transgenic sorghum events, the higher absolute bioaccessibility findings indicate these lines would have potential for public health impact to consumers of traditional porridges through enhanced dietary delivery of provitamin and non-provitamin A carotenoids as well as mineral elements.

Mineral deficiencies including iron and zinc are a major problem in sub-Saharan Africa and disproportionately affect women and children (Black et al., 2008; Harika et al., 2017). It has been previously estimated that addressing zinc deficiencies could prevent 4.4% of childhood deaths (Fischer Walker et al., 2009). Biofortification strategies have sought to improve mineral content in crops through enhanced uptake and/or partitioning in edible tissues (Khush et al., 2012; Díaz-Gómez et al., 2017; Debelo et al., 2020). However, mineral content is poorly associated with bioavailability (Glahn & Noh, 2021). Mineral elements co-localize in sorghum tissues (e.g. pericarp) with phytate and bioavailability is largely influenced by this anti-nutrient due to its ability to chelate metal ions (Ferruzzi et al., 2020).

We explored the effects of activating both naturally occurring and heterologous phytase enzymes on the profiles of inositol phosphates as well as carotenoid bioaccessibility. Phytase has long been known to catabolize phytate into various inositol phosphate products in legumes (Frias et al., 2003) as well as cereals including sorghum, corn, rice, and wheat (Azeke et al., 2011). Similar trends were observed in the current study as phytase activation shifted inositol phosphate profiles, normally dominated by phytate (IP6), to smaller metabolites (IP1-5; Fig. 3, Table S4). Notably, the heterologous PhyA in our transgenic events not only functioned more efficiently than the naturally present enzyme, but also reduced total inositol phosphate pools in sorghum flours. This finding was evident in significant shifts in calculated molar ratios of phytate to zinc and iron (Fig. 4). Lower molar ratios of phytate to zinc and iron are known to be correlated with mineral bioaccessibility (Morris & Ellis, 1989; Ma et al., 2007). While estimates vary, critical values for molar phytate to zinc or iron are generally believed to fall between 10-15 (Oberleas & Harland, 1981; Turnlund et al., 1984; Morris & Ellis, 1989; Saha et al., 1994). Above these thresholds, bioaccessibility is impaired due to the chelating activity of phytate. Our study demonstrated that as a result of phytase activation, transgenic materials exhibited the most drastic reduction of phytate to iron ratio from 16.0 to 4.1 and phytate to zinc ratio from 19.0 to 4.9 (Fig. 4), far below reported critical thresholds. However, with additional iron and zinc liberated and available for absorption, a potential challenge emerges due to the known antagonistic relationship between carotenoids and minerals.

Previous studies have shown that divalent cations in concentrations above certain thresholds are able to precipitate bile salts (Gu et al., 1992) as well as fatty acids (Atteh & Leeson, 1985). The resulting formation of insoluble soaps has been hypothesized to be a potential mechanism by which carotenoid bioaccessibility is negatively impacted by minerals (Graham & Sackman, 1983; Biehler et al., 2011a; Corte-Real et al., 2016, 2018). In this study, total phytate remaining in steeped, transgenic events was on average 22.9% of the original amount and additional minerals were ostensibly released through phytate degradation. However, no significant effect was observed on relative or absolute carotenoid bioaccessibility in our study as a function of pre-steeping and phytase activation. This outcome may be due to the inherently low concentrations of mineral elements in sorghum relative to the treatments tested in other bodies of work in which concentrations of minerals tended to fall within the mmol range (Graham & Sackman, 1983; Biehler et al., 2011a; Corte-Real et al., 2016, 2018; Kopec et al., 2019). When prepared as a traditional porridge, an average 200 g serving of porridge (40 g of flour) from all germplasm studied in this work contains 1.62 and 1.47 mg (29.01 and 22.48 µmol/L) of iron and zinc, respectively. Given the effects of divalent minerals on micellarization are concentration dependent (Biehler et al., 2011a; Corte-Real et al., 2016, 2018), we hypothesize that the low mineral content of sorghum may obscured any significant effects of steeping, phytase activation, and subsequent liberation of minerals on micellarization efficiency. Critical values above which micellarization is negatively affected for divalent cations such as zinc, magnesium, and calcium have previously been reported at 100, 300, and 500 mg/L, respectively (Corte-Real et al., 2017). While iron and zinc both fell below 40 mg/L in the sorghum material studied here, magnesium was present at 1584.4 mg/L on average. The relatively higher concentrations of magnesium likely contributed to modest reductions observed. However, the modest decreases in micellarization efficiency observed in steeped transgenic sorghum porridges were not statistically significant. This finding is consistent with studies conducted by our group in other crops such as spinach (Hayes et al., 2021). Regardless, the quantity of carotenoids available for absorption were not significantly affected by steeping and the subsequent release of divalent cations. Should future iterations of this material yield transgenic events capable of delivering larger quantities of minerals per serving, it would be prudent to ensure micellarization efficiency and deliverable carotenoids are not significantly impacted. Given the chemical aspects of the sorghum germplasm in this study, our findings indicate that the transgenic events reported here can effectively deliver both carotenoids and divalent minerals simultaneously and more efficiently than non-transgenic controls.

## 1.4.1 Conclusions

The present study sought to leverage biotechnology for the purpose of generating nutritionally enhanced cultivars of sorghum. We utilized these events to study interactions between divalent minerals and carotenoid bioaccessibility, as well as the role of naturally occurring and heterologous phytase on these processes. Data generated in this study indicate that transgenic events reported here can simultaneously deliver significant quantities of provitamin A carotenoids, carotenoids associated with eye and brain development, and divalent minerals required for normal growth and development. Importantly, increasing the deliverability of mineral elements by enzymatically degrading phytate did not significantly compromise carotenoid bioavailability. While human trials are needed to determine the clinical efficacy of these sorghum events, our data suggest that simultaneously addressing multiple (and competing) nutrient deficiencies is feasible and warrants additional attention.

## Supporting information

Collection of supplemental tables

## 1.5 Acknowledgements

We thank Sydney Corbin, Jacob Fitzgerald, Micaela Hayes, Zulfiqar Mohamedshah, and Candace Nunn for their assistance during *in vitro* digestion and carotenoid extraction experiments. This work was supported by a gift from the Pioneer Foundation (to MGF) and in part by the U.S. Department of Agriculture, Agricultural Research Service (**USDA-ARS Project 6026-51000-012** and **3092-51000-061-000)**

## 1.7 Official capacity disclaimer

The findings and conclusions in this publication are those of the authors and should not be construed to represent any official USDA or U.S. Government determination or policy. Mention of trade names or commercial products in this publication is solely for the purpose of providing specific information and does not imply recommendation or endorsement by the U.S. Department of Agriculture. The USDA is an equal opportunity provider and employer.

## 1.8 Material availability

Novel biological materials described in this publication may be available to the academic community and other not-for-profit institutions solely for non-commercial research purposes upon acceptance and signing of a material transfer agreement between the author’s institution and the requestor. In some cases, such materials may contain genetic elements described in the manuscript that were obtained from a third party(s) *(e*.*g*., *Zm-WS1 whole seed promoter (Fu et al*., *2010))* and the authors may not be able to provide materials including third party genetic elements to the requestor because of certain third-party contractual restrictions placed on the author’s institution. In such cases, the requester will be required to obtain such materials directly from the third party. The author’s and authors’ institution do not make any express or implied permission(s) to the requester to make, use, sell, offer for sale, or import third party proprietary materials. Obtaining any such permission(s) will be the sole responsibility of the requestor. Corteva Agriscience™ proprietary plant germplasm and any transgenic material will not be made available except at the discretion of the owner and then only in accordance with all applicable governmental regulations.

## Works Cited

Abbitt, S.E., Jung, R., & Selinger, D.A. (2009). Seed-Preferred Promoters. US20090049571A1

Afify, A.E.-M.M.R., El-Beltagi, H.S., El-Salam, S.M.A., & Omran, A.A. (2011). Bioavailability of Iron, Zinc, Phytate and Phytase Activity during Soaking and Germination of White Sorghum Varieties. PLOS ONE, 6, e25512. https://doi.org/10.1371/journal.pone.0025512

Albertsen, M.C., Anderson, P.C., Che, P., Glassman, K.F., Jung, R., & Zhao, Z.-Y. (2014). Transformed plants having increased beta-carotene levels, increased half-life and bioavailability and methods of producing such. WO2014070646A1

Aragón, I.J., Ceballos, H., Dufour, D., & Ferruzzi, M.G. (2018). Pro-vitamin A carotenoids stability and bioaccessibility from elite selection of biofortified cassava roots (Manihot esculenta, Crantz) processed to traditional flours and porridges. Food & Function, 9, 4822–4835. https://doi.org/10.1039/c8fo01276h

Atteh, J.O., & Leeson, S. (1985). Influence of Age, Dietary Cholic Acid, and Calcium Levels on Performance, Utilization of Free Fatty Acids, and Bone Mineralization in Broilers. Poultry Science, 64, 1959–1971. https://doi.org/10.3382/ps.0641959

Azeke, M.A., Egielewa, S.J., Eigbogbo, M.U., & Ihimire, I.G. (2011). Effect of germination on the phytase activity, phytate and total phosphorus contents of rice (Oryza sativa), maize (Zea mays), millet (Panicum miliaceum), sorghum (Sorghum bicolor) and wheat (Triticum aestivum). Journal of Food Science and Technology, 48, 724–729. https://doi.org/10.1007/s13197-010-0186-y

Biehler, E., Hoffmann, L., Krause, E., & Bohn, T. (2011)(a). Divalent Minerals Decrease Micellarization and Uptake of Carotenoids and Digestion Products into Caco-2 Cells. The Journal of Nutrition, 141, 1769–1776. https://doi.org/10.3945/jn.111.143388

Biehler, E., Kaulmann, A., Hoffmann, L., Krause, E., & Bohn, T. (2011)(b). Dietary and host-related factors influencing carotenoid bioaccessibility from spinach (Spinacia oleracea). Food Chemistry, 125, 1328–1334. https://doi.org/10.1016/j.foodchem.2010.09.110

Black, R.E., Allen, L.H., Bhutta, Z.A., Caulfield, L.E., de Onis, M., Ezzati, M., Mathers, C., & Rivera, J. (2008). Maternal and child undernutrition: global and regional exposures and health consequences. The Lancet, 371, 243–260. https://doi.org/10.1016/S0140-6736(07)61690-0

Cahoon, E.B., Hall, S.E., Ripp, K.G., Ganzke, T.S., Hitz, W.D., & Coughlan, S.J. (2003). Metabolic redesign of vitamin E biosynthesis in plants for tocotrienol production and increased antioxidant content. Nature Biotechnology, 21, 1082–1087. https://doi.org/10.1038/nbt853

Che, P., Anand, A., Wu, E., Sander, J.D., Simon, M.K., Zhu, W., Sigmund, A.L., Zastrow-Hayes, G., Miller, M., Liu, D., Lawit, S.J., Zhao, Z.-Y., Albertsen, M.C., & Jones, T.J. (2018). Developing a flexible, high-efficiency Agrobacterium-mediated sorghum transformation system with broad application. Plant Biotechnology Journal, 16, 1388–1395. https://doi.org/10.1111/pbi.12879

Che, P., Zhao, Z.-Y., Glassman, K., Dolde, D., Hu, T.X., Jones, T.J., Gruis, D.F., Obukosia, S., Wambugu, F., & Albertsen, M.C. (2016). Elevated vitamin E content improves all-trans β-carotene accumulation and stability in biofortified sorghum. Proceedings of the National Academy of Sciences, 113, 11040–11045. https://doi.org/10.1073/pnas.1605689113

Che, P., Zhao, Z.-Y., Hinds, M., Rinehart, K., Glassman, K., & Albertsen, M. (2019). Evaluation of Agronomic Performance of β-Carotene Elevated Sorghum in Confined Field Conditions. In Z.-Y. Zhao & J. Dahlberg (Eds.), Sorghum: Methods and Protocols (pp. 209–220). Springer, New York, NY.

Corte-Real, J., Bertucci, M., Soukoulis, C., Desmarchelier, C., Borel, P., Richling, E., Hoffmann, L., & Bohn, T. (2017). Negative effects of divalent mineral cations on the bioaccessibility of carotenoids from plant food matrices and related physical properties of gastro-intestinal fluids. Food & Function, 8, 1008–1019. https://doi.org/10.1039/C6FO01708H

Corte-Real, J., Desmarchelier, C., Borel, P., Richling, E., Hoffmann, L., & Bohn, T. (2018). Magnesium affects spinach carotenoid bioaccessibility in vitro depending on intestinal bile and pancreatic enzyme concentrations. Food Chemistry, 239, 751–759. https://doi.org/10.1016/j.foodchem.2017.06.147

Corte-Real, J., Iddir, M., Soukoulis, C., Richling, E., Hoffmann, L., & Bohn, T. (2016). Effect of divalent minerals on the bioaccessibility of pure carotenoids and on physical properties of gastro-intestinal fluids. Food Chemistry, 197, 546–553. https://doi.org/10.1016/j.foodchem.2015.10.075

Coruzzi, G., Broglie, R., Cashmore, A., & Chua, N.H. (1983). Nucleotide sequences of two pea cDNA clones encoding the small subunit of ribulose 1,5-bisphosphate carboxylase and the major chlorophyll a/b-binding thylakoid polypeptide.. Journal of Biological Chemistry, 258, 1399–1402. https://doi.org/10.1016/S0021-9258(18)32995-8

Cunningham, F.X., & Gantt, E. (1998). Genes and Enzymes of Carotenoid Biosynthesis in Plants. Annual Review of Plant Physiology and Plant Molecular Biology, 49, 557–583. https://doi.org/10.1146/annurev.arplant.49.1.557

Da, S., Akingbala, J.O., Rooney, L., Scheuring, J.F., & Miller, F.R. (1981). In L.W. Rooney & D.S. Murty (Eds.), Evaluation of Tô Quality in a Sorghum Breeding Program. (pp. 11–23). ; International Crops Research Institute for the SemiArid Tropics, Patancheru, A.P., India.

Debelo, H., Albertsen, M., Simon, M., Che, P., & Ferruzzi, M. (2020). Identification and Characterization of Carotenoids, Vitamin E and Minerals of Biofortified Sorghum. Current Developments in Nutrition, 4, 1792. https://doi.org/10.1093/cdn/nzaa067_019

Díaz-Gómez, J., Twyman, R.M., Zhu, C., Farré, G., Serrano, J.C., Portero-Otin, M., Muñoz, P., Sandmann, G., Capell, T., & Christou, P. (2017). Biofortification of crops with nutrients: factors affecting utilization and storage. Current Opinion in Biotechnology, 44, 115–123. https://doi.org/10.1016/j.copbio.2016.12.002

Failla, M.L., Chitchumroonchokchai, C., Siritunga, D., De Moura, F.F., Fregene, M., Manary, M.J., & Sayre, R.T. (2012). Retention during processing and bioaccessibility of β-carotene in high β-carotene transgenic cassava root. Journal of Agricultural and Food Chemistry, 60, 3861–3866. https://doi.org/10.1021/jf204958w

Ferruzzi, M.G., Kruger, J., Mohamedshah, Z., Debelo, H., & Taylor, J.R.N. (2020). Insights from in vitro exploration of factors influencing iron, zinc and provitamin A carotenoid bioaccessibility and intestinal absorption from cereals. Journal of Cereal Science, 96, 103126. https://doi.org/10.1016/j.jcs.2020.103126

Fischer Walker, C.L., Ezzati, M., & Black, R.E. (2009). Global and regional child mortality and burden of disease attributable to zinc deficiency. European Journal of Clinical Nutrition, 63, 591–597. https://doi.org/10.1038/ejcn.2008.9

Fraval, S., Hammond, J., Bogard, J.R., Ng’endo, M., van Etten, J., Herrero, M., Oosting, S.J., de Boer, I.J.M., Lannerstad, M., Teufel, N., Lamanna, C., Rosenstock, T.S., Pagella, T., Vanlauwe, B., Dontsop-Nguezet, P.M., Baines, D., Carpena, P., Njingulula, P., Okafor, C., Wichern, J., Ayantunde, A., Bosire, C., Chesterman, S., Kihoro, E., Rao, E.J.O., Skirrow, T., Steinke, J., Stirling, C.M., Yameogo, V., & van Wijk, M.T. (2019). Food Access Deficiencies in Sub-saharan Africa: Prevalence and Implications for Agricultural Interventions. Frontiers in Sustainable Food Systems, 3

Frias, J., Doblado, R., Antezana, J.R., & Vidal-Valverde, C. (2003). Inositol phosphate degradation by the action of phytase enzyme in legume seeds. Food Chemistry, 81, 233–239. https://doi.org/10.1016/S0308-8146(02)00417-X

Fu, H., Brown, J.A., Francis, K., & Song, H.-S. (2010). Whole seed specific promoter. WO2010122110A1

Glahn, R.P., & Noh, H. (2021). Redefining Bean Iron Biofortification: A Review of the Evidence for Moving to a High Fe Bioavailability Approach. Frontiers in Sustainable Food Systems, 5

Graham, D.Y., & Sackman, J.W. (1983). Solubility of calcium soaps of long-chain fatty acids in simulated intestinal environment. Digestive Diseases and Sciences, 28, 733–736. https://doi.org/10.1007/BF01312564

Gu, J., Hofmann, A., Ton-Nu, H., Schteingart, C., & Mysels, K. (1992). Solubility of calcium salts of unconjugated and conjugated natural bile acids.. Journal of Lipid Research, 33, 635–646. https://doi.org/10.1016/S0022-2275(20)41428-2

Han, Y., Wilson, D.B., & Lei, X.G. (1999). Expression of an Aspergillus niger phytase gene (phyA) in Saccharomyces cerevisiae. Applied and Environmental Microbiology, 65, 1915–1918. https://doi.org/10.1128/AEM.65.5.1915-1918.1999

Harika, R., Faber, M., Samuel, F., Kimiywe, J., Mulugeta, A., & Eilander, A. (2017). Micronutrient Status and Dietary Intake of Iron, Vitamin A, Iodine, Folate and Zinc in Women of Reproductive Age and Pregnant Women in Ethiopia, Kenya, Nigeria and South Africa: A Systematic Review of Data from 2005 to 2015. Nutrients, 9, 1096. https://doi.org/10.3390/nu9101096

Hayes, M., Corbin, S., Nunn, C., Pottorff, M., Kay, C.D., Lila, M.A., Iorrizo, M., & Ferruzzi, M.G. (2021). Influence of simulated food and oral processing on carotenoid and chlorophyll in vitro bioaccessibility among six spinach genotypes. Food & Function, 12, 7001–7016. https://doi.org/10.1039/D1FO00600B

Hayes, M., Pottorff, M., Kay, C., Van Deynze, A., Osorio-Marin, J., Lila, M.A., Iorrizo, M., & Ferruzzi, M.G. (2020). In Vitro Bioaccessibility of Carotenoids and Chlorophylls in a Diverse Collection of Spinach Accessions and Commercial Cultivars. Journal of Agricultural and Food Chemistry, 68, 3495–3505. https://doi.org/10.1021/acs.jafc.0c00158

Humphrey, J.H., West, K.P., & Sommer, A. (1992). Vitamin A deficiency and attributable mortality among under-5-year-olds.. Bulletin of the World Health Organization, 70, 225–232

Institute of Medicine (US) Committee on Nutrition Standards for National School Lunch and Breakfast Programs. (2008). Nutrition Standards and Meal Requirements for National School Lunch and Breakfast Programs: Phase I. Proposed Approach for Recommending Revisions. In V.A. Stallings & C.L. Taylor (Eds.), National Academies Press (US), Washington (DC).

Kayodé, A.P., Linnemann, A.R., Nout, M.J., & Van Boekel, M.A. (2007). Impact of sorghum processing on phytate, phenolic compounds and in vitro solubility of iron and zinc in thick porridges. Journal of the Science of Food and Agriculture, 87, 832–838. https://doi.org/10.1002/jsfa.2782

Kayodé, A.P.P., Linnemann, A.R., Hounhouigan, J.D., Nout, M.J.R., & van Boekel, M.A.J.S. (2006). Genetic and Environmental Impact on Iron, Zinc, and Phytate in Food Sorghum Grown in Benin. Journal of Agricultural and Food Chemistry, 54, 256–262. https://doi.org/10.1021/jf0521404

Kean, E.G., Bordenave, N., Ejeta, G., Hamaker, B.R., & Ferruzzi, M.G. (2011). Carotenoid bioaccessibility from whole grain and decorticated yellow endosperm sorghum porridge. Journal of Cereal Science, 54, 450–459. https://doi.org/10.1016/j.jcs.2011.08.010

Khush, G.S., Lee, S., Cho, J.-I., & Jeon, J.-S. (2012). Biofortification of crops for reducing malnutrition. Plant Biotechnology Reports, 6, 195–202. https://doi.org/10.1007/s11816-012-0216-5

Kopec, R.E., Caris-Veyrat, C., Nowicki, M., Bernard, J.-P., Morange, S., Chitchumroonchokchai, C., Gleize, B., & Borel, P. (2019). The Effect of an Iron Supplement on Lycopene Metabolism and Absorption During Digestion in Healthy Humans. Molecular Nutrition & Food Research, 63, 1900644. https://doi.org/10.1002/mnfr.201900644

Kruger, J., Oelofse, A., & Taylor, J.R.N. (2014). Effects of aqueous soaking on the phytate and mineral contents and phytate:mineral ratios of wholegrain normal sorghum and maize and low phytate sorghum. International Journal of Food Sciences and Nutrition, 65, 539–546. https://doi.org/10.3109/09637486.2014.886182

Li, S., Nugroho, A., Rocheford, T., & White, W.S. (2010). Vitamin A equivalence of the β-carotene in β-carotene–biofortified maize porridge consumed by women. The American Journal of Clinical Nutrition, 92, 1105–1112. https://doi.org/10.3945/ajcn.2010.29802

Lipkie, T.E., De Moura, F.F., Zhao, Z.-Y., Albertsen, M.C., Che, P., Glassman, K., & Ferruzzi, M.G. (2013). Bioaccessibility of Carotenoids from Transgenic Provitamin A Biofortified Sorghum. Journal of Agricultural and Food Chemistry, 61, 5764–5771. https://doi.org/10.1021/jf305361s

Ma, G., Li, Y., Jin, Y., Zhai, F., Kok, F.J., & Yang, X. (2007). Phytate intake and molar ratios of phytate to zinc, iron and calcium in the diets of people in China. European Journal of Clinical Nutrition, 61, 368–374. https://doi.org/10.1038/sj.ejcn.1602513

Mares, J. (2016). Lutein and Zeaxanthin Isomers in Eye Health and Disease. Annual Review of Nutrition, 36, 571–602. https://doi.org/10.1146/annurev-nutr-071715-051110

Mendiburu, F. de, & Yanseen, M. (2021). agricolae: Statistical Procedures for Agricultural Research

Morris, E.R., & Ellis, R. (1989). Usefulness of the dietary phytic acid/zinc molar ratio as an index of zinc bioavailability to rats and humans. Biological Trace Element Research, 19, 107–117. https://doi.org/10.1007/BF02925452

Negrotto, D., Jolley, M., Beer, S., Wenck, A.R., & Hansen, G. (2000). The use of phosphomannose-isomerase as a selectable marker to recover transgenic maize plants (Zea mays L.) via Agrobacterium transformation. Plant Cell Reports, 19, 798–803. https://doi.org/10.1007/s002999900187

Neudert, U., Martínez-Férez, I.M., Fraser, P.D., & Sandmann, G. (1998). Expression of an active phytoene synthase from Erwinia uredovora and biochemical properties of the enzyme. Biochimica Et Biophysica Acta, 1392, 51–58. https://doi.org/10.1016/s0005-2760(98)00017-4

Oberleas, D., & Harland, B.E. (1981). Phytate content of foods: Effect on dietary zinc bioavailability. Journal of the American Dietetic Association, 79, 433–436. https://doi.org/10.1016/S0002-8223(21)39390-7

Paine, J.A., Shipton, C.A., Chaggar, S., Howells, R.M., Kennedy, M.J., Vernon, G., Wright, S.Y., Hinchliffe, E., Adams, J.L., Silverstone, A.L., & Drake, R. (2005). Improving the nutritional value of Golden Rice through increased pro-vitamin A content. Nature Biotechnology, 23, 482–487. https://doi.org/10.1038/nbt1082

Park, S., Ong, R.G., & Sticklen, M. (2016). Strategies for the production of cell wall[deconstructing enzymes in lignocellulosic biomass and their utilization for biofuel production. Plant Biotechnology Journal, 14, 1329–1344. https://doi.org/10.1111/pbi.12505

Peng, Y., Schertz, K.F., Cartinhour, S., & Hart, G.E. (1999). Comparative genome mapping of Sorghum bicolor (L.) Moench using an RFLP map constructed in a population of recombinant inbred lines. Plant Breeding, 118, 225–235. https://doi.org/10.1046/j.1439-0523.1999.118003225.x

R Development Core Team. (2018). R: A language and environment for statistical computing.

Ram, K., Wickham, H., Richards, C., & Baggett, A. (2018). wesanderson: A Wes Anderson Palette Generator

Saha, P.R., Weaver, C.M., & Mason, A.C. (1994). Mineral Bioavailability in Rats from Intrinsically Labeled Whole Wheat Flour of Various Phytate Levels. Journal of Agricultural and Food Chemistry, 42, 2531–2535. https://doi.org/10.1021/jf00047a029

Tang, G., Qin, J., Dolnikowski, G.G., Russell, R.M., & Grusak, M.A. (2009). Golden Rice is an effective source of vitamin A. The American Journal of Clinical Nutrition, 89, 1776–1783. https://doi.org/10.3945/ajcn.2008.27119

Thomas, S.E., & Johnson, E.J. (2018). Xanthophylls. Advances in Nutrition, 9, 160–162. https://doi.org/10.1093/advances/nmx005

Trumbo, P., Yates, A.A., Schlicker, S., & Poos, M. (2001). Dietary Reference Intakes: Vitamin A, Vitamin K, Arsenic, Boron, Chromium, Copper, Iodine, Iron, Manganese, Molybdenum, Nickel, Silicon, Vanadium, and Zinc. Journal of the American Dietetic Association, 101, 294–301. https://doi.org/10.1016/S0002-8223(01)00078-5

Turnlund, J.R., King, J.C., Keyes, W.R., Gong, B., & Michel, M.C. (1984). A stable isotope study of zinc absorption in young men: effects of phytate and a-cellulose. The American Journal of Clinical Nutrition, 40, 1071–1077. https://doi.org/10.1093/ajcn/40.5.1071

Weber, D., & Grune, T. (2012). The contribution of β-carotene to vitamin A supply of humans. Molecular Nutrition & Food Research, 56, 251–258. https://doi.org/10.1002/mnfr.201100230

Whitcher, J.P., Srinivasan, M., & Upadhyay, M.P. (2001). Corneal blindness: a global perspective. Bulletin of the World Health Organization, 79, 214–221. https://doi.org/10.1590/S0042-96862001000300009

Wickham, H. (2016). Ggplot2 Elegant Graphics for Data Analysis (Use R!). Springer Springer-Verlag, New York, NY.

Xu, M., Luan, Y., Zhang, Z., & Jiang, S. (2021). Dietary pattern changes over Africa and its implication for land requirements for food. Mitigation and Adaptation Strategies for Global Change, 26, 13. https://doi.org/10.1007/s11027-021-09939-4

Ye, X., Al-Babili, S., Klöti, A., Zhang, J., Lucca, P., Beyer, P., & Potrykus, I. (2000). Engineering the provitamin A (beta-carotene) biosynthetic pathway into (carotenoid-free) rice endosperm. Science (New York, N.Y.), 287, 303–305. https://doi.org/10.1126/science.287.5451.303

You, H. (2016). Quantifying the bioefficacy of β-carotene-biofortified sorghum [Doctor of Philosophy]. Iowa State University, Digital Repository, Ames

Zhang, S., Yang, W., Zhao, Q., Zhou, X., Fan, Y., & Chen, R. (2017). Rapid Method for Simultaneous Determination of Inositol Phosphates by IPC-ESI–MS/MS and Its Application in Nutrition and Genetic Research. Chromatographia, 2, 275–286. https://doi.org/10.1007/s10337-017-3238-x

